# The Effects of Strain and Estrous Cycle on Heroin- and Sucrose-Maintained Responding in Female Rats

**DOI:** 10.1101/2020.11.06.370023

**Authors:** Karl T. Schmidt, Jessica L. Sharp, Sarah B. Ethridge, Tallia Pearson, Shannon Ballard, Kenzie M. Potter, Mark A. Smith

## Abstract

Heroin intake decreases during the proestrus phase of the estrous cycle in female, Long-Evans rats. The purpose of this study was to (1) determine if proestrus-induced decreases in heroin intake extend across rat strains and (2) determine if proestrus-induced decreases in responding extend to a nondrug reinforcer. Female rats were implanted with intravenous catheters and trained to self-administer heroin. Estrous cycle was tracked daily for the duration of the study. During testing, Lewis, Sprague-Dawley, and Long Evans rats self-administered low (0.0025 mg/kg) and high (0.0075 mg /kg) doses of heroin (Experiment 1) and then self-administered sucrose (Experiment 2) on fixed ratio (FR1) schedules of reinforcement. Heroin intake decreased significantly during proestrus in all three rat strains under at least one dose condition; however, sucrose intake did not decrease during proestrus in any strain. These data indicate that responding maintained by heroin, but not a nondrug reinforcer, significantly decreases during proestrus in female rats and that these effects are consistent across rat strain.

## 1. Introduction

Previously, our laboratory reported a decrease in heroin self-administration during the proestrus phase of the female estrous cycle (Lacy et al., 2016, Smith et al., 2020a; Smith et al., 2020b). Consistently, we observed a reduction in heroin intake between 55% and 70% compared to sessions during other phases of the estrous cycle. However, one caveat is that each of these studies was conducted using Long-Evans rats, and it is unknown whether this effect is unique to this strain of rats. Furthermore, previous examinations of the effects of estrous cycle on heroin self-administration have neglected examinations of changes in responding maintained by a nondrug reinforcer. The possibility remains that the reported decrease in self-administration during proestrus is a result of non-specific motoric effects or a proestrus-induced change in motivated behavior. Therefore, the purpose of this study was to determine whether the effects of proestrus on heroin-self administration extend to other commonly used laboratory rat strains and are specific to the heroin stimulus. We measured heroin intake across the estrous cycle in female Lewis (LEW), Sprague Dawley (SD), and Long-Evans (LE) rats. Following heroin testing, rats were tested to determine levels of sucrose-maintained responding across the estrous cycle.

## 2. Methods

### 2.1 Animals

Three strains of female rats were obtained from Charles River Laboratories (Raleigh, NC): LEW, SD, and LE. Rats arrived at 49 days old and were housed individually. Rats were maintained on a 12-hr light/dark cycle (lights on: 0500-1700) with ad libitum access to food and water, except during initial lever-press training. All husbandry and procedures were approved by the Animal Care and Use Committee of Davidson College.

### 2.2 Experimental Chambers

Heroin self-administration took place in custom built, operant conditioning chambers (Faircloth Machine Shop, Winston-Salem, NC) that also served as home cages (Lacy et al., 2014; Smith, 2012). These chambers allowed rats to respond on a single response lever inserted into the chamber during test sessions. Responses on the lever activated a drug infusion via a syringe pump outside the chamber. Med Associates software (St. Albans, VT) recorded response data and interfaced between response levers and pump activation.

Lever-press training and sucrose self-administration occurred in separate, commercially available operant conditioning chambers (Med Associates Inc, St. Albans, VT). Chambers included two response levers, stimulus lights above the response levers, a food hopper between the response levers capable of delivering either 45 mg grain pellets (BioServ, Flemington NJ) or 45 mg sucrose pellets (BioServ), and a house light opposite the response levers.

### 2.3 Lever-Press Training

Rats were food restricted to approximately 90% of their free-feeding body weight beginning one week following arrival. Rats were trained to lever press using food reinforcement during daily 2 hr sessions. In each session, responses were reinforced on a continuous reinforcement (FR1) schedule during which each lever press delivered a grain pellet. Rats were determined to be trained once 40 reinforcers were earned in four sessions. Once rats were trained to lever press, ad libitum food was available for the remainder of the study.

### 2.4 Surgery

Once trained to lever press, rats were anesthetized with an intraperitoneal injection of 100 mg/kg ketamine and 8 mg/kg xylazine. Indwelling jugular catheters were implanted as described previously (Smith et al., 2008). Postoperatively, rats were treated with topical antibiotic ointment and subcutaneous injections of 3 mg/kg ketoprofen for 2 days. Intravenous antibiotic and heparinized saline were infused through the catheter daily for 7 days to prevent infection and maintain catheter patency. After one week, heparinized saline was administered twice per week to maintain catheter patency.

### 2.5 Heroin Self-Administration

Approximately one week after surgery, rats were transferred to heroin self-administration chambers that served as home cages during drug self-administration testing. At this time, estrous phase monitoring began as described previously (Lacy et al., 2016; Smith et al., 2020b). Briefly, approximately one hour prior to the start of each self-administration session, samples of vaginal cells were obtained via vaginal lavage and examined under light microscope to classify as metestrus, diestrus, proestrus, or estrus. Across multiple studies (Lacy et al., 2016; Smith et al., 2020a; Smith et al., 2020b), we have seen no differences between metestrus and diestrus on heroin intake, so data from these phases were combined for analysis.

Self-administration sessions began with the insertion of the lever into the chamber and a noncontingent infusion of heroin. Responses were reinforced on an FR1 schedule by an infusion of 0.0075 mg/kg heroin. Each infusion was followed by a 20 s timeout during which the lever was retracted. Sessions lasted two hours and terminated with the retraction of the lever. Self-administration training continued in this fashion for one complete estrous cycle, which was typically 4-6 days. After one complete estrous cycle was completed, rats were randomized to receive either 0.0075 mg/kg/inf or 0.0025 mg/kg/inf. All experimental conditions during testing were identical to those used during training. Testing continued at a particular dose until vaginal cytology provided a clear indication that each phase of the estrous cycle was represented. If multiple cycles were required to obtain data from each phase, data from repeated phases were averaged for each subject. After obtaining data for each phase of the estrous cycle, the other dose of heroin was tested. Again, daily sessions continued until each phase of the estrous cycle was represented.

### 2.6 Sucrose Self-Administration

Following heroin self-administration, rats were transferred to individual polycarbonate cages for the remainder of the study. Daily one-hour test sessions occurred in operant conditioning chambers separate from those used during heroin self-administration. During these sessions, each lever press was reinforced with a sucrose pellet on an FR1 schedule. Estrous cycle phase was monitored approximately one hour prior to test sessions as described above. For each individual rat, daily sessions continued until the rat was tested during each phase of the estrous cycle. If multiple cycles were required to obtain data from each phase, data from repeated phases were averaged for each subject.

### 2.7 Data Analysis

Data obtained during heroin self-administration sessions were operationalized as number of reinforcers (i.e., number of infusions, number of lever presses) per session. The effects of strain were determined by collapsing the data across estrous phase and analyzing via ANOVA, followed by post hoc tests using Tukey’s honestly significant difference (HSD) correction for multiple comparisons. The effects of estrous phase were determined via repeated-measures ANOVA for each strain and dose of heroin. In cases in which the omnibus test was significant, estrous phases were compared via paired-samples t-tests using the Holms-Bonferroni correction for multiple comparisons. Effects sizes were characterized using the partial eta squared statistic (η^2^) using standard definitions of effect size (small: 0.01; medium: 0.09; large: 0.25). Data obtained in the sucrose self-administration tests were operationalized as number of reinforcers (i.e., number of sucrose pellets, number of lever presses) per session and analyzed in an identical fashion. All statistical tests were two-tailed and used an alpha level of .05.

## 3. Results

Heroin self-administration was readily acquired in all rats (data not shown), but intake varied across strain and phase of estrous. Intake of 0.0075 mg/kg heroin varied significantly across strain (*F*(2, 38) = 9.696; *p* < .001), with a large effect size (η^2^ = .338) and rank order of SD > LE ≈ LEW (Figure 1A). Responding varied as a function of estrous phase in both LE (*F*(2, 24) = 13.912; *p* < .001) and LEW (*F*(2, 20) = 15.071; *p* < .0011) rats, but not SD rats. Within the LEW and LE strains, responding was significantly lower during proestrus than during met/diestrus and estrus. The effect sizes were large and comparable in both LE (η^2^ = .537) and LEW (η^2^ = .601) strains.

**Figure 1.**
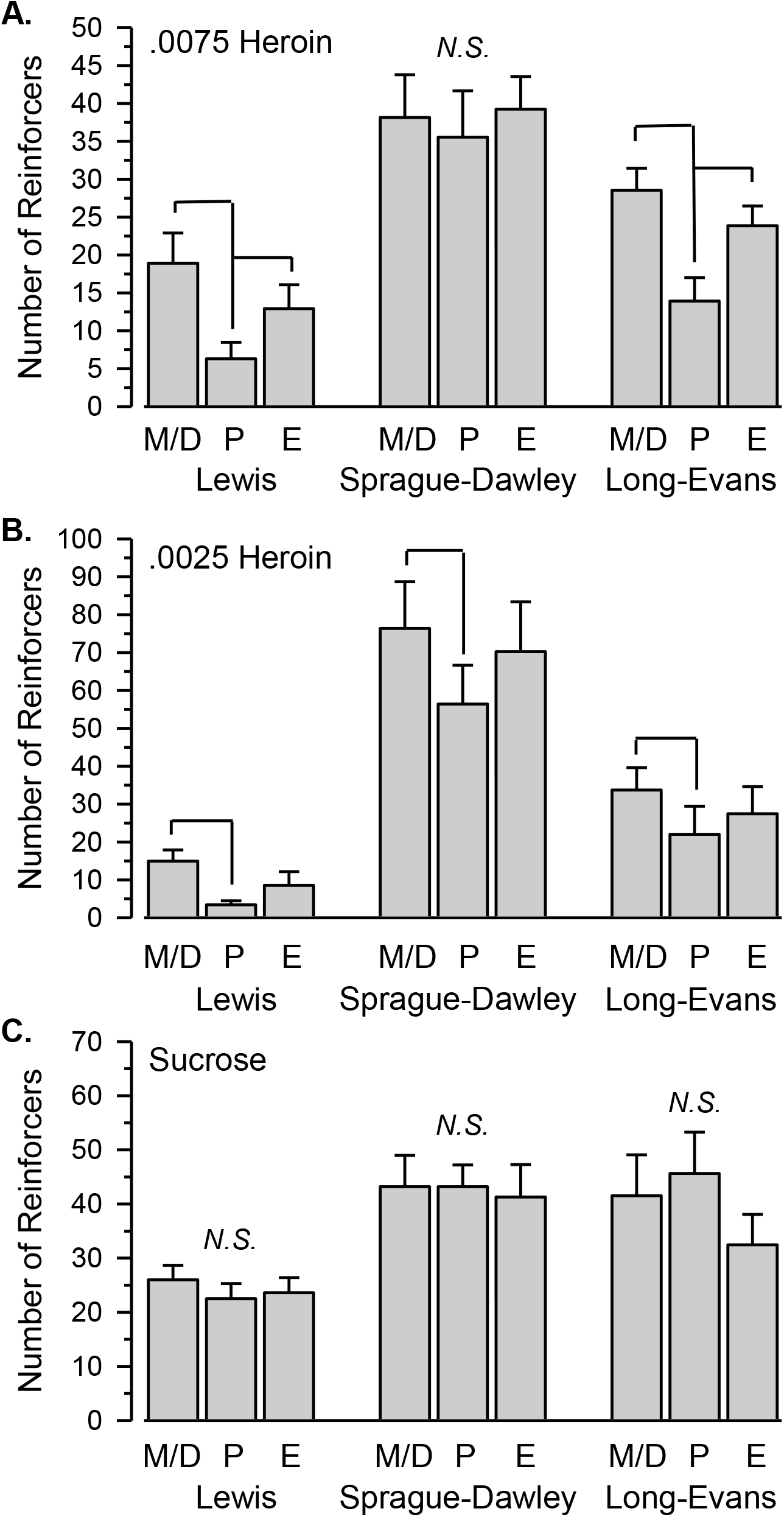
Number of reinforcers during 2-hr test sessions in female rats responding during met/diestrus (M/D), proestrus (P), and estrus (E) phases of the estrous cycle. Figure 1A depicts mean (SE) number of reinforcers maintained by .0075 m/kg/infusion heroin in Lewis (n = 11), Sprague-Dawley (n = 17), and Long-Evans (n = 13) rats. Figure 1B depicts mean (SE) number of reinforcers maintained by .0025 m/kg/infusion heroin in Lewis (n = 5), Sprague-Dawley (n = 13), and Long-Evans (n = 11) rats. Figure 1C depicts mean (SE) number of reinforcers maintained by 45 mg sucrose pellets in Lewis (n = 16), Sprague-Dawley (n = 15), and Long-Evans (n = 14) rats. Horizontal linkage lines indicate significant differences between phases of the estrous cycle. N.S. indicates no significant main effect of estrous cycle.

Similar to that observed with the higher dose of heroin, intake of 0.0025 mg/kg heroin varied significantly across strain (*F*(2, 26) = 8.575; *p* = .001), with a large effect size (η^2^ = .397) and rank order of SD > LE ≈ LEW (Figure 1B). Responding varied as a function of estrous phase in all three rats strains (SD: *F*(2, 24) = 4.912; *p* = .016; LE: *F*(2, 20) = 3.620; *p* = .046; LEW: *F*(2, 8) = 5.028; *p* = .039). In all three strains, responding was significantly lower during proestrus than met/diestrus. Responding was numerically less during proestrus than estrus in all three strains, but in no strain were these differences statistically significant after correcting for multiple comparisons. The effect sizes were large in all strains, but markedly greater in LEW (η^2^ = .557) than both SD (η^2^ = .290) and LE (η^2^ = .266) strains.

Sucrose self-administration was acquired in all rats during the first training session despite the move to new operant conditioning chambers. Responding maintained by sucrose differed significantly across strains (*F*(2, 41) = 5.589; *p* = .007), with a moderate effect size (η^2^ = .214) and rank order of SD ≈ LE > LEW (Figure 1C). Unlike that observed with heroin self-administration, responding did not differ across the estrous cycle in any strain.

## 4. Discussion

Here, we show that proestrus-induced decreases in heroin self-administration can be observed across three commonly used strains of laboratory rats and that, in all three strains, this decrease in responding does not extend to responding maintained by sucrose. These data contribute to a growing literature investigating changes in drug self-administration as a function of ovarian hormones.

Early investigations examining the effects of ovarian hormones (on opioid self-administration reported mixed results, with one study reporting estradiol increasing the rate of acquisition of heroin self-administration (Roth et al., 2002) and one study reporting that estradiol does not influence heroin self-administration (Stewart et al., 1996). However, more recent studies have reported that estradiol reduces heroin seeking across multiple experimental conditions (Sedki et al., 2015; Vazquez et al., 2020). Results from our laboratory are consistent with these most recent studies, given that we have shown that opioid self-administration is reduced during proestrus in an estrogen but not progesterone dependent manner (Lacy et al., 2016; Smith et al., 2020a; Smith et al., 2020b). Similar processes are likely responsible for the reduction in heroin-maintained responding in the present study.

Previous demonstrations of proestrus-induced decreases in heroin intake were conducted in LE rats, and it was unknown whether these effects were idiosyncratic to this strain. One purpose of the present study was to determine if proestrus-induced decreases in heroin intake are observed across different strains of rats commonly used in preclinical research. LE rats were chosen to maintain consistency with previous studies (e.g., Lacy et al., 2016; Smith et al., 2020a; Smith et al., 2020b) and served as the representative comparison group. LEW rats were chosen because they are more sensitive to the positive reinforcing effects of opioids relative to other strains, and because they have been described as a possible genetic model of drug abuse vulnerability (Cadoni, 2016). Lastly, SD rats were chosen because they are the most common strain used in self-administration studies, appearing in nearly double the number of studies than the closest alternative strain (determined via PubMed search performed on October 12, 2020). In the present study, proestrus-induced decreases in heroin self-administration were observed in each strain under at least one dose condition, despite significant strain differences in overall levels of heroin intake. These findings indicate that proestrus-induced decreases in heroin intake are generalizable across three rats strains that differ in baseline levels of responding. This latter observation is also consistent with a previous report that heroin intake decreases during proestrus across individual rats regardless of whether the rat is a high or low responder (Lacy et al., 2016).

One additional purpose of this study was to determine whether proestrus-induced decreases in responding extend to a non-drug reinforcer. Analysis of responding by a nondrug reinforcer allows a determination of whether decreases in heroin-maintained responding are specific to the heroin stimulus or due to nonspecific effects on motoric or motivated behavior. Responding maintained by sucrose did not vary across the estrous cycle in any of the three strains tested, despite significant strain differences in overall sucrose intake. Consequently, the effects of proestrus on heroin self-administration cannot be attributed to factors related to nonspecific motoric or motivational processes.

## 5. Conclusions

In conclusion, this study replicates the finding that heroin intake decreases significantly during the proestrus phase of the estrous cycle in normally cycling female rats. Importantly, the study demonstrates that this effect is apparent across three strains of laboratory rats, despite significant strain differences in overall heroin intake. Also of importance, proestrus-induced decreases in responding does not extend to a nondrug reinforcer, indicating that the effect is not due to proestrus-induced decreases in motoric or motivated behavior. Collectively, these data reveal that heroin intake, but not sucrose intake, is influenced by circulating concentrations of endogenous ovarian hormones. We are not aware of any studies that have examined the influence of ovarian hormones on measures of opioid intake or opioid seeking in human populations; however, these data suggest that manipulations of ovarian hormones may serve as a target for future pharmacotherapies for women with opioid use disorder.

## Acknowledgements

The authors thank the National Institute on Drug Abuse for supplying the study drug.

